# Germline genomic patterns are associated with cancer risk, oncogenic pathways and clinical outcomes

**DOI:** 10.1101/616268

**Authors:** Xiaowen Feng, Xue Xu, Derek Li, Qinghua Cui, Edwin Wang

**Author notes:** Correspondence (Qinghua Cui), (Edwin Wang).

## Abstract

Germline genetic polymorphism is prevalent and inheritable. So far mutations of a handful of genes have been associated with cancer risks. For example, women who harbor *BRCA1/2* germline mutations have a 70% of cumulative breast cancer risk; individuals with congenital germline *APC* mutations have nearly 100% of cumulative colon cancer by the age of fifty. At present, gene-centered cancer predisposition knowledge explains only a small fraction of the inheritable cancer cases. Here we conducted a systematic analysis of the germline genomes of cancer patients (n=9,712) representing 22 common cancer types along with non-cancer individuals (n=16,670), and showed that seven germline genomic patterns, or significantly repeatedly occurring sequential mutation profiles, could be associated with both carcinogenesis processes and cancer clinical outcomes. One of the genomic patterns was significantly enriched in the germline genomes of patients who smoked than in those of non-smoker patients of 13 common cancer types, suggesting that the germline genomic pattern was likely to confer an elevated carcinogenesis sensitivity to tobacco smoke. Several patterns were also associated with somatic mutations of key oncogenic genes and somatic-mutational signatures which are associated with higher genome instability in tumors. Furthermore, subgroups defined by the germline genomic patterns were significantly associated with distinct oncogenic pathways, tumor histological subtypes and prognosis in 12 common cancer types, suggesting that germline genomic patterns enable to inform treatment and clinical outcomes. These results demonstrated that genetic cancer risk and clinical outcomes could be encoded in germline genomes in the form of not only mutated genes, but also specific germline genomic patterns, which provided a novel perspective for further investigation.

## Introduction

Germline genetic polymorphism is prevalent and inheritable. The contribution of genetic predisposition to cancer development and progression has been recognized for centuries and has yet been widely investigated(Friedman and Iwai, 1976),(McPherson, 2000),(Nakagawa et al., 2004). Compared to somatic mutations in tumors, germline malignant variants face looser selection pressure and are inherited along with numerous passenger mutations(Bignell et al., 2010). Systematic genome sequencing of normal tissues of cancer patients provides a considerable chance to study the germline genomic variants.

As the cancer-driving genetic germline variants distribute sparsely across genomes and restricted to a small set of genes(Bignell et al., 2010), investigations of germline genetic variants have been largely restricted to the known cancer driver genes including tumor suppressors and the ones closely related to DNA repair, oncogenic signaling pathways, cell cycle and so on(Farmer et al., 2005),(Rahman, 2014). For example, individuals diagnosed as colorectal cancer among the first-degree relatives who have Lynch syndrome(VASEN et al., 1999) have been shown to carry DNA repair defects that disabled DNA damage resurrection in germlines and therefore accumulated genetic alterations which led to colon cancer. Germline mutated Ras has been shown to be associated with developmental disorders(Schubbert et al., 2006) and cardio-facio-cutaneous(Niihori et al., 2006). Germline *BRCA1/2* mutations have been well-known to be directly associated with increased risks in multiple cancer types including breast and ovarian cancer(Gayther et al., 1999),(Antoniou et al., 2003),(Maxwell et al., 2014).

Although gene-centered studies have proven to be informative, so far only a handful of associations between genes and cancer risk have been determined(Fletcher and Houlston, 2010). Thus, we asked if a genomic pattern, or a significantly repeatedly occurring sequential profile in germline genomes, could serve as a novel measurement for malignant genetic predisposition. To answer this question, we conducted a systematic analysis of the germline genomes of 9,712 cancer patients representing 22 cancer types and 16,670 non-cancer individuals and revealed seven cancer-related germline genomic patterns. One of them (i.e., susceptibility genomic pattern for smoking, or a conditional-genetic risk factor) was significantly more enriched in the germlines of the smoker patients than in those of non-smokers patients. These results suggested that germline genomic patterns could provide an inspiring and novel measurement for cancer susceptibility. Furthermore, germline subgroups defined by the patterns were significantly correlated to clinically meaningful differences in terms of cancer histological subtypes, oncogenic mechanisms and survival outcomes in 12 common cancer types. These results implied that cancer genetic risk could be encoded in germline genomes in the form of not only genes such as *BRCA1/2* but also genomic patterns. We speculated that further analysis of germline genomic patterns may unravel genetic mechanisms of diseases where molecular mechanisms could be beyond the current gene-centered paradigm.

## Results

### Genomic patterns in cancer germline genomes

We obtained 430,772,708 germline substitutions from the whole-exome sequencing data of 9,712 cancer patients in The Cancer Genome Atlas (TCGA)(Grossman et al., 2016), representing 22 cancer types, and 46,998,783 somatic substitutions from their paired tumor genomes. Germline mutational catalog was generated by summarizing potential substitution profiles, where variations were incorporated with sequential sequence-contexts. Meanwhile, whole-exome data of 16,670 non-cancer patients of three cohorts (see Methods) were merged to form a non-cancer dataset, served as the background. Blood cancers were not included for further analysis (see Methods).

Genetic variants aggregated in a germline genome could be viewed as accumulated outcomes of mutational processes that happened to its ancestral genomes and evolutionary genetic polymorphisms. A germline genomic pattern presents a pan-genome enrichment of recurring substitutions in a sequence-context within germline genomes. Intuitively, a genomic pattern pertaining to cancer risk is most likely to display detectable enrichment in the germline genomes of cancer patients. Genomic patterns are buried in the high-dimensional sequence features in genomes so that extracting the genomic patterns becomes a tedious and computationally intensive task by applying statistical methods. Non-negative matrix factorization (NMF) enables interpretable feature extraction from high-dimensional data(Berry et al., 2007), (Lee and Seung, 2001) and has been employed in extracting of biological meaningful somatic mutational signatures in tumors(Nik-Zainal et al., 2012), (Alexandrov et al., 2013b), (Alexandrov et al., 2013a). Therefore, we adapted this approach for germline mutational catalogs (Method; Supplementary Materials Figure S1 and S2), demonstrated its stability and then applied it to the germline mutational catalogs of the cancer patients and non-cancer population (for background noise reduction), respectively. By doing so, we identified seven distinct genomic patterns (named cancer-associated germline genomic pattern, CGGP) (Figure 1; Supplementary Materials Data D1) along with their contributions which were represented by weighting factors, respectively. Further analysis confirmed that none of the genomic patterns were sequencing artifacts(Nik-Zainal et al., 2012).

**Figure 1.**
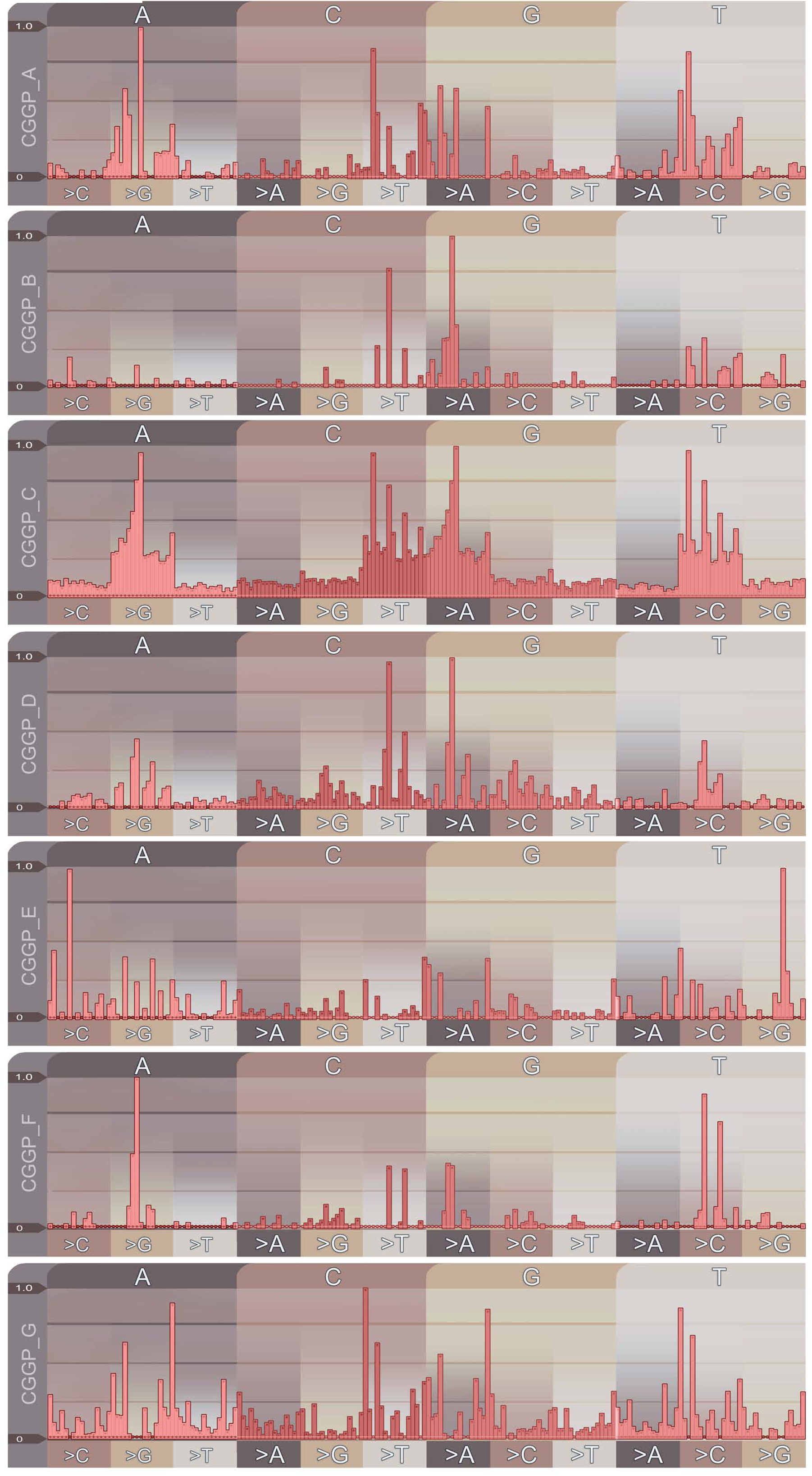
Cancer-associated germline genomic patterns deciphered from the germline genomes of cancer patients. Components of the genomic patterns are displayed according to their substitutions (reference: upper label; mutation: lower label) and immediate 5’ and 3’ sequential contexts. Signal strengths displayed at the vertical bars indicate the overall prevalence of the patterns.

Figure 1 illustrated the visualization of seven CGGPs. CGGP_A, D, and F were mainly characterized by an enrichment of transitional mutations. CGGP_F was characterized by A>G and T>C variants, while CGGP_A and CGGP_D contained mainly C/T>T/C and G/A>A/G variants. Other context-dependent mutations were also revealed in these three patterns, and when combined with other CGGPs, CGGP_A and CGGP_D mainly contributed to lung, pancreas and stomach cancers.

Signals of C<u>G</u>G>C<u>A</u>G and C<u>C</u>G>C<u>T</u>G were the traits of CGGP_B. When utilized in combinations with CGGP_C or CGGP_E, CGGP_B was a potential contributor to the brain, breast, cervix, colon, and rectum, kidney, lung and stomach cancers (see below).

CGGP_C appeared to be a dominant pattern, with the strongest assigned signals and weights in samples amongst CGGPs. This pattern was mainly accounted for the transitions of A>G, G>A, C>T and T>C with modest preferences for sequence contexts.

Conspicuous signals in CGGP_E led to the formation of a local triple nucleotide profile of repeated nucleotides, including C<u>A</u>C>C<u>C</u>C, G<u>A</u>G>G<u>G</u>G, A<u>C</u>A>A<u>A</u>A, G<u>C</u>G>G<u>G</u>G, C<u>G</u>C>C<u>C</u>C, T<u>G</u>T>T<u>T</u>T, C<u>T</u>C>C<u>C</u>C and G<u>T</u>G>G<u>G</u>G. We found that CGGP_E was a genetic susceptibility germline genomic pattern for tobacco smoke, and the smokers whose germline carried CGGP_E had significantly elevated risks in 13 common cancer types. Although enrichment of CGGP_E would imply biologically meaningful information in certain cancer types, the combination of multiple CGGPs provides much more insights (see below).

CGGP_G was characterized in the preference of T<u>A/G</u>T>T<u>G/A</u>T and A<u>C/T</u>A>A<u>T/C</u>A mutations as well as modest overall assigned signals in other mutational profiles. Weights (i.e., representing the contribution of the pattern to a sample) for CGGP_G in germline genomes of cancer patients were significantly higher than the weights granted by non-cancer individuals, the differences between whom were also the most prominent in seven CGGPs (Supplementary Materials Figure. S3).

To examine whether the seven genomic patterns were cancer-specific, we applied our modified NMF approach to the germline exome sequences of non-cancer individuals (n=16,670) and identified 6 genomic patterns. Patterns except CGGP_E were reproduced (cosine similarities were: 0.004, 0.023, 0.0005, 0.026, 0.072 and 0.007 for CGGP_ A, B, C, D and F, respectively). We further extended this analysis to the germline dataset derived from the combination of the cancer patients and non-cancer individuals (n=9,712+16,670), and found that all 7 patterns were reproducible. These results implied that while CGGP_E might be a pattern that’s significantly more enriched in cancer patients’ germlines than in non-cancer individuals.

### CGGP_E was a susceptibility genomic pattern for tobacco smoke

A genetic predisposition could collaborate with exogenous cancer risk factors to drive tumorigenesis. While tobacco smoke is a well-recognized exogenous risk factor for inducing cancer, less than 15% of the smokers would end up developing lung cancer. It is evidential that individual fates are not only affected by exposing to mutagens but also their genetically determined sensitivity to mutagens. Thus, we asked if any CGGP could be associated with tobacco smoking in cancer patients. We, therefore, examined the enrichment of each CGGP between the germlines of smoker and non-smoker patients.

Signatures 4 and 29 of somatic mutational signatures have been reported in the Catalogue Of Somatic Mutations In Cancer (COSMIC) (http://cancer.sanger.ac.uk/cosmic/signatures) to be the somatic mutational imprints of tobacco smoking and tobacco chewing habit found in tumors. It has been reported that at least 17 cancer types were linked to smoking(Alexandrov et al., 2016). We partitioned the cancer patients into two groups: one was affected by tobacco smoking and the other was not, based on the contribution of Signatures 4 and 29 in tumors using the methods described previously (Alexandrov et al., 2013a)). Comparative analyses of each of the CGGPs in the germline genomes between smoking and non-smoking groups revealed that CGGP_E was significantly enriched in the smoking group across 13 common cancer types, including lung (n=974, t-statistic=24.9, p-value=1.05×10^-101^, two-tailed t-test), prostate (n=494, t=33.3, p=3.4×10^-119^), bladder (n=405, t=20.5, p=2.35×10^−61^), head and neck (n=503, t=26.6, p=1.0×10^−90^), kidney (n=674, t=17.4, p=3.9×10^−55^), stomach (n=433, t=19.6, p=4.3×10^−59^), thyroid (n=486, p=16.9, t=5.1×10^−49^), colorectal (n=578, t=3.3, p=3.0×10^−3^), breast (n=971, t=8.6, p=1.18×10^−15^), liver (n=367, t=4.1, p=3.7×10^−5^) and uterus cancers (n=564, t=2.7, p=6.0×10^−3^) (Figure 2A). Cancer types with insufficient sample sizes were not examined. A higher weight of CGGP_E in germline genome had a significant positive correlation with the presence of smoking-related somatic mutational signature (COSMIC Signature 4) in tumor genome as for whole patient population (Figure 2B). These results suggested that CGGP_E could be a susceptibility CGGP for smoking, i.e., smoking alone could not necessarily drive tumorigenesis unless the CGGP_E was presented in the germline genome of a tobacco smoker. Furthermore, in general, higher contribution of CGGP_E in samples implied shorter survivals (p=0.01, log-rank test). More significant results were observed in four cancer types including adrenal gland (1.1×10^−4^), bladder (p=5.3×10^−4^), kidney (p=6.2×10^−3^), and stomach (p=3.3×10^−3^) cancers. These results suggested that CGGP_E could have a quantitative effect on cancer development and prognosis. Surprisingly, such a correlation was not observed in lung cancer (see Discussion). Further, we obtained similar results when assigning CGGP_E in a binary fashion (see Methods) to patients.

**Figure 2.**
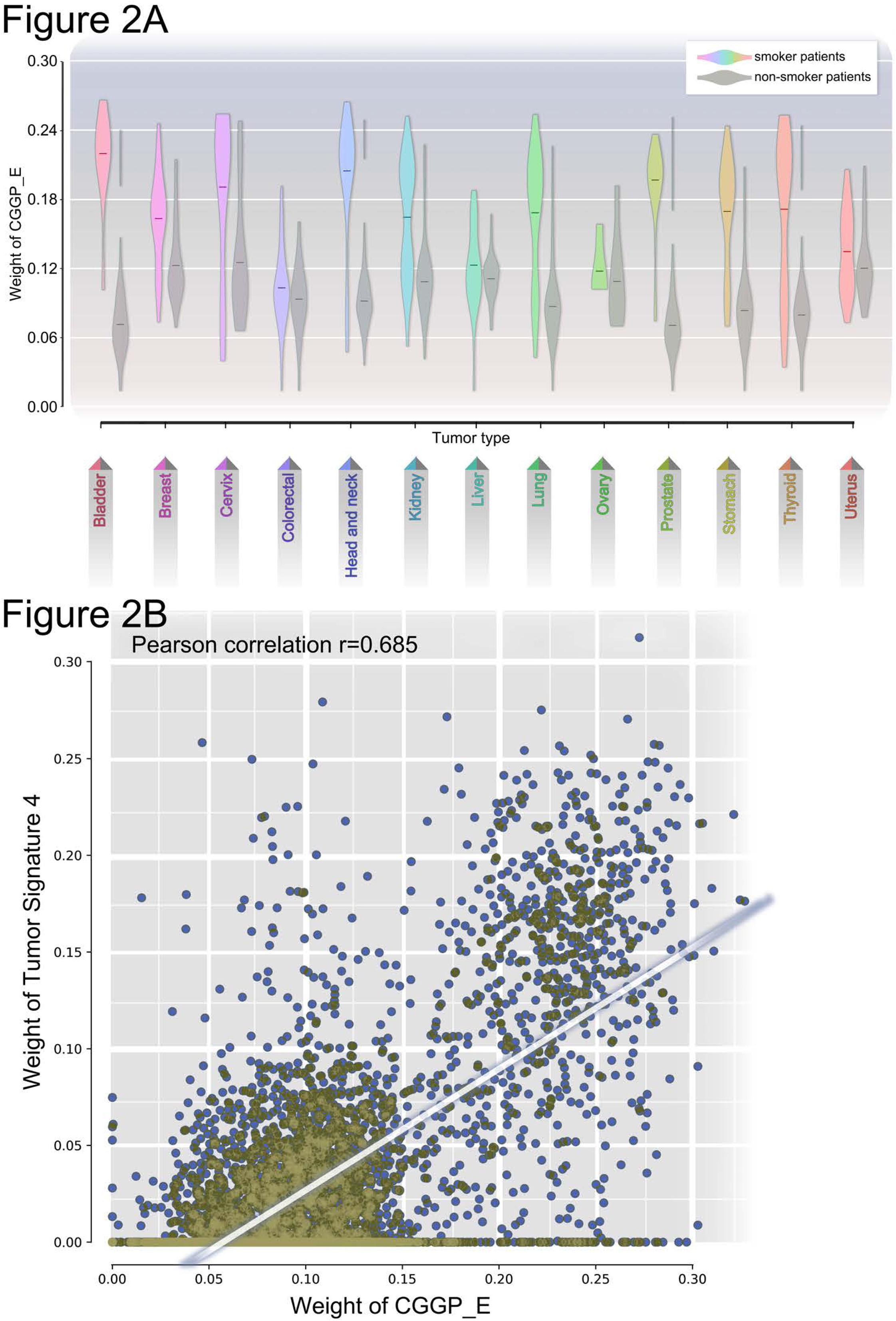
The relationship between germline genomic patterns and tobacco mutagen sensitivity. **(A)** Germline genomic pattern (CGGP_E) is highly enriched in the smoker patients than the non-smoker patients in 13 cancer types (p<10^−5^, t-test). Grey violin-boxes represent non-smoking patients, while the boxes with other colors represent smoking patients. **(B)** Contribution (i.e., weight) of CGGP_E in germline genomes was significantly correlated to the possession of smoking-related somatic mutational signatures in their paired tumors.

To further understand the molecular mechanisms of CGGP_E’s impact on cancer development, we ranked patients based on the contributions of CGGP_E in germlines in descending order and selected the patients from the upper quartile and the lower quartile to form two groups. Genes containing mutations that fitted CGGP_E were examined and compared between the two groups. Well-known cancer drivers, such as fibronectin type-III (*FN3*), receptors of tyrosine kinases (*RTKs*), EGF-like domain proteins and insulin-like growth factor binding proteins, were significantly more frequently mutated in the upper quartile group (FDR-corrected p<0.05; Supplementary Materials Data D2). These results suggested that the presence of CGGP_E in germlines might introduce significantly more mutations to *RTKs* and other cancer driver genes, which in return could induce higher carcinogenesis risks for the individuals when exposing to mutagenic agents. Thus, we reasoned that except tobacco smoke the CGGP_E might be able to be a susceptibility genomic pattern for other mutagens. This hypothesis could be further tested in the future when related data become available.

These results provided a potential rationale for the long-established observation that less than 20% of heavy smokers would develop lung cancer in their lifetime. When extending the same analysis to other CGGPs for understanding of molecular mechanisms, we did not obtain biologically meaningful results. However, in specific tumor types, other CGGPs showed a casual positive correlation with the smoking signatures (e.g., CGGP_G was significantly enriched in smoker patients of colorectal (p=3.2×10^−7^), head and neck (p=3.6×10^−5^), kidney (p=9.1×10^−18^), lung (p=9.3×10^−19^), prostate (p=1.0×10^−32^), stomach (p=2.9×10^−15^) and thyroid (p=6.4×10^−6^) cancer groups), suggesting latent mechanisms might still to be revealed for the CGGPs.

### CGGPs impacted on somatic mutation of key oncogenic genes in tumors

To explore whether the CGGPs were associated with somatic mutations of key cancer drivers and family history of cancer, in each cancer type we partitioned the patients into two subgroups, either carried any mutation in the given gene, or did not, and examined whether the weighing factors of CGGPs differed between the subgroups (see Methods). Due to the sample size limitation, only most frequently somatically mutated genes across all cancer types reported by TCGA were examined: TP53, PIK3CA, KMT2D, FAT4, ARID1A, PTEN, KMT2C, APC, KRAS, FAT1, ATRX, NF1, ZFHX3, IDH1, ATM, TRRAP, RNF213, AKAP9, GRIN2A. Patients of each cancer type were partitioned based on whether a given gene was nonsynonymously mutated in their tumor tissues, and distributions of CGGP weighing factors were examine between such subgroups.

Our observations implied that the germline patterns had impacts on which somatic mutated genes were to be selected by tumors (Figure 3, Supplementary Materials Table S1). In 19 cancer types we observed that enrichment of certain CGGPs were positively correlated with higher somatical mutation frequency of the above genes (see Methods). Across all cancer types, TP53, PTEN, and FAT1 mutations were associated with higher weighing of CGGP_B or D (p=8.1×10^−3^ and 1.7×10^− 2^), CGGP_A (p=1.0×10^−2^), and CGGP_E (p=4.0×10^−3^), respectively. In individual cancer types, such correlations were observed between EGFR somatic mutations and CGGP_F in lung cancer (p=1.37×10^−5^), and between CGGP_F and IDH1 somatic mutation (p=2.7×10^−44^) and a significantly higher probability of having a family history of cancers (p=1.1×10^−14^) in brain cancer population.

**Figure 3.**
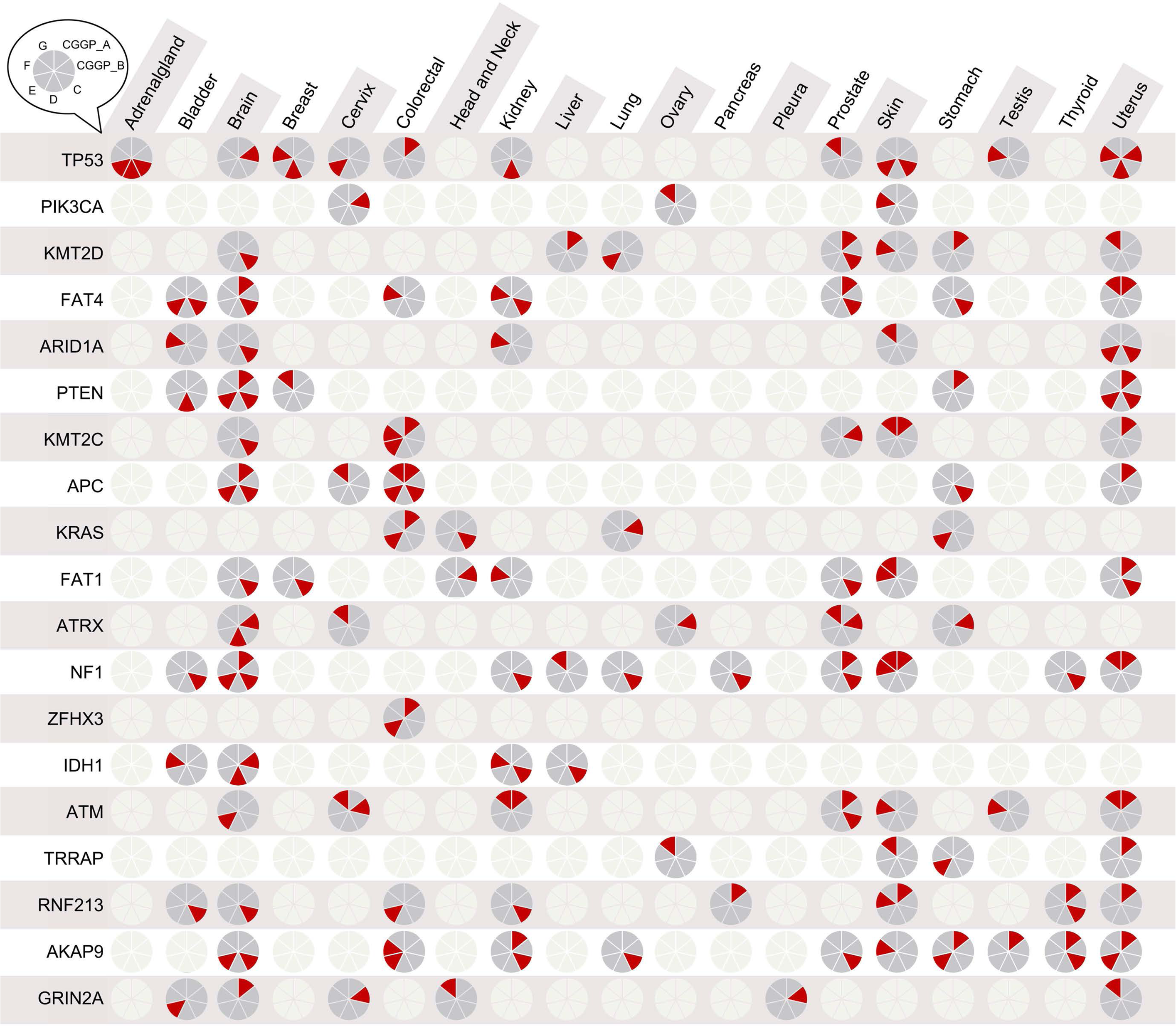
Enrichment of CGGPs implied somatic mutations in ontogenetic genes in tumor tissue. Significantly elevated weighing factors of CGGPs were found correlated with abundancy of ontogenetic mutations in somatic genomes of 19 cancer types, 19 most frequently mutated genes across all cancer types. Each small pie chart in the plot had either one color (ivory) or two colors (gray and red), where the former one indicated no significant result was observed in the corresponding cancer type & somatic gene mutation pair, and the later one showed which CGGP(s) was(were) implying the somatic gene mutation with its red slots (starting from the top right slot and going clockwise, CGGP_A to CGGP_G alphabetically; p<0.05; see Methods).

These results suggested that germline genomic patterns exert evo-constrains on somatic mutated genes in tumors, and though CGGP_E was prominent in cancer patients, the combinations of multiple germline genomic patterns might exert stronger evo-constrains on selecting of tumor somatic mutations. Therefore, CGGP combinations could serve as features for classifying cancer patients through thresholding or unsupervised clustering methods.

### CGGP combinations in germlines were associated with cancer types

In the search for meaningful CGGP combinations, we conducted statistical analyses of systematic combinations of the CGGPs (k=2, 3). Germline data of non-cancer population (n=16,670) was utilized to justify background noise, i.e. we considered a CGGP to be closely linked to cancer risks only when it contributed to the carrier at a significantly higher level than it did amongst the non-cancer population. Randomization tests (10,000 iterations) demonstrated that combination of the CGGPs were significantly more prevalent in cancer patients compared to non-cancer population, and germline groups defined by CGGP combinations (k=2 and 3) were significantly enriched in distinct cancer types (Figure 4). Few patients had more than three CGGPs in their germline genomes. These results have an implication of germline pattern’s role in tissue-specific carcinogenesis. For example, patients with higher contributions of CGGP_E and CGGP_G in their germlines were more enriched in being diagnosed with brain, kidney, lung and head and neck cancers (p=4.5×10^−23^, 3.0×10^−31^, 5.0×10^−8^, 6.6×10^−6^, ratio=2.3, 3.0,1.6,1.7, Fisher’s test; n=1,464). Colorectal and prostate cancer diagnoses were significantly enriched in the patients (n=90) with CGGP_D and CGGP_G (p=9.2×10^−17^, 0.04, ratio=9.7, 2.1, Fisher’s test). Along with this line, the combination of CGGP_B, CGGP_C and CGGP_G was enriched in ovary cancer group (p=1.3×10^− 23^, ratio=27.4). Furthermore, patients carried the combination of CGGP_D, CGGP_E and CGGP_G in their germlines were significantly enriched 10 COSMIC somatic mutational signatures in their tumors tissues compared to the patients without the combination (p=1.1×10^−2^, 3.4×10^−2^, 3.0×10^−2^, 6.6×10^−5^, 3.6×10^−6^, 1.4×10^−2^, 6.4×10^−11^, 2.9×10^−3^, 1.1×10^−8^ and 7.8×10^−3^, respectively; Supplementary Materials Table S2), implying germline genomic patterns had the potential to shape or select somatic mutational processes during tumorigenesis. Tumors of the patients who carry the combination of CGGP_C, CGGP_D and CGGP_E were enriched only with COSMIC Signature 14 in their paired tumors. COSMIC Signature 14 has been annotated to be associated with elevated somatic mutation burden as high as 200 mutations per MB in tumors, suggesting that the combination of CGGP_C, CGGP_D and CGGP_E in germlines could imply a higher genome instability. The patients carrying CGGP_E and CGGP_G had an enrichment of COSMIC Signature 3 and 6 in their tumors. COSMIC has annotated that Signature 3 and 6 are strongly associated with elevated number of long or short insertions and deletions, respectively. Interestingly, COSMIC also reported a strong association between Signature 3 and germline BRCA1/2 mutations in three cancer types as well. In summary, these results suggested that CGGPs or their combinations could be associated with triggering of endogenous mutations in tumors, and furthermore, could possibly shape the carcinogenesis and tumor proliferation process.

**Figure 4.**
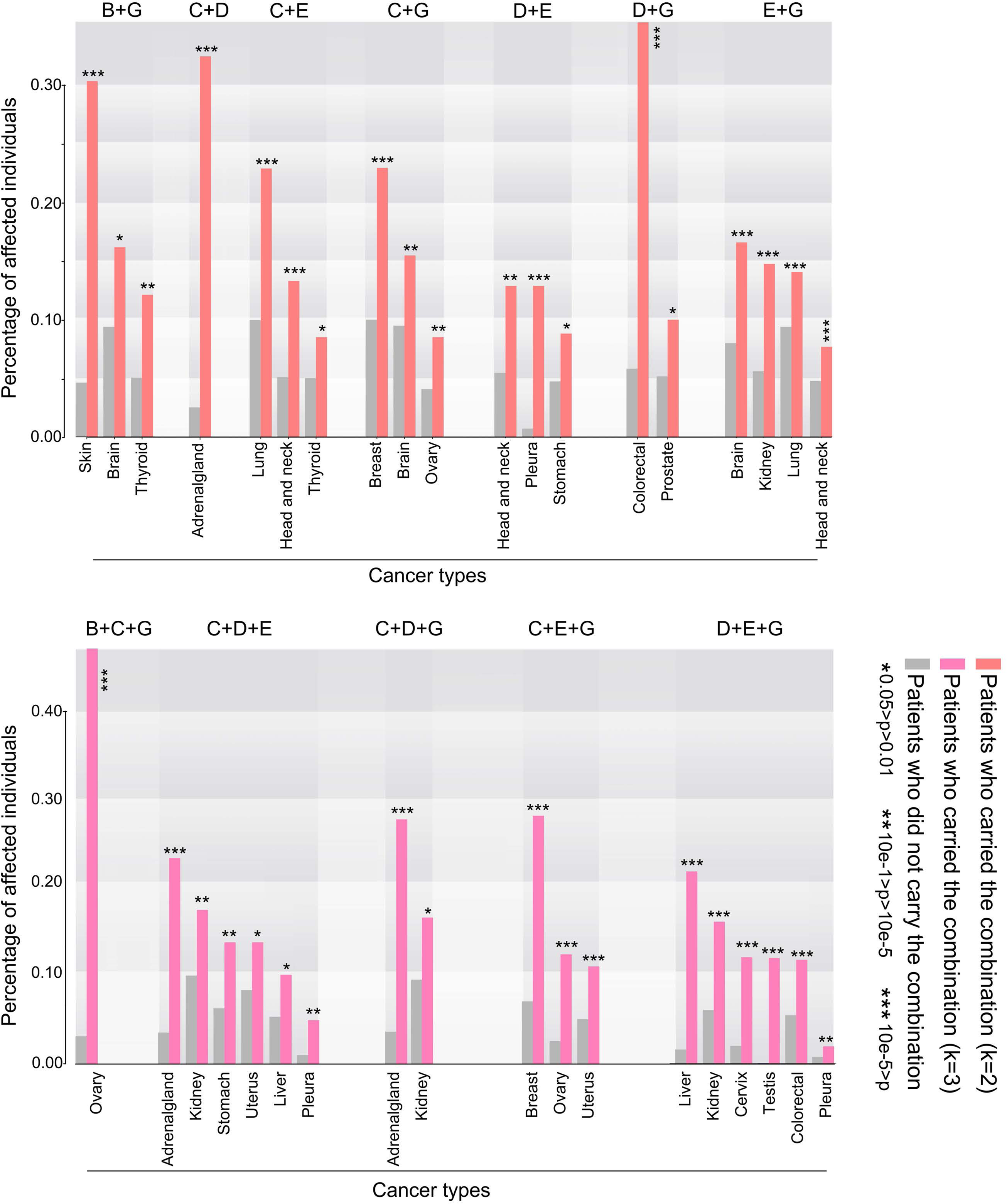
combinations of cancer-associated germline genomic patterns were associated with cancer types. For each tumor type, patients were divided into two groups (carrier group and non-carrier group) based on whether they carried a given combination of CGGPs. CGGP combinations demonstrated in the bar plots were enriched in several tumor types for their carrier, implying that CGGP combinations could be related to specific tumorigenesis process. CGGP combinations that had less than 50 samples in either subgroups were excluded. For example, skin, brain and thyroid cancers were significantly enriched in patients who carried the combination of CGGP_B and CGGP_G in germline genomes (Supplementary Materials Table S2).

### CGGP-defined germline subgroups were associated with distinct tumor histological subtypes, oncogenic pathways and prognosis

We next explored if CGGPs could classify germline genomes into subgroups for a given cancer type. To do so, we clustered the patients of each cancer type (i.e., cancer types with insufficient sample sizes were excluded for the analysis) based on CGGPs-contribution profiles in germlines. The partitions were conducted using the unsupervised hierarchical clustering. For most cancer types, the CGGPs were able to classify germlines into stable subgroups; by applying dimension reduction methods such as PCA ahead of hierarchical clustering we obtained similar results). CGGPs-defined germline subgroups exhibited correlations with cancer histological subtypes in four cancer types: brain, lung, kidney and stomach cancers. Cancer types lacked histology diagnosis information were not analyzed. Brain cancer patients were clustered into three germline subgroups. One subgroup (subgroup 1, beige in Figure 5A) was enriched with glioblastoma multiform (GBM) samples, while subgroup 3 (subgroup 3, gray in Figure 5A) was significantly enriched with less-aggressive astrocytoma samples. Subgroup 2 was also enriched with GBM samples but, unlike subgroup 1, was in favor of CGGP-A rather than CGGP_E (subgroup 1+2 n=322, subgroup 2 n=575, p=1.2×10^−26^ for GBM, χ2 test). GBM−enriched germline subgroups had significantly shorter survival than the other subgroup (p=2.2×10^−31^ and 3.1×10^−26^). Likewise, three germline subgroups were found in kidney cancer patients. Two of the subgroups were enriched with clear cell renal carcinoma, the other was dominated by kidney papillary renal cell carcinoma and kidney chromophobe cancer (p=1.0×10^−5^, χ2 test). Significant survival differences were also observed between the subgroups (subgroup 1 vs 2: p=1.9×10^−2^, subgroup 1 vs 3: p=6.3×10^−4^, sugroup2 vs 3: p=5.6×10^−4^). Furthermore, two germline subgroups of lung cancer patients were significantly enriched with adenocarcinoma and squamous cell carcinoma samples, respectively (n=179 and 795, p=1.1×10^−2^). These results suggested that CGGPs in the germline genomes of these three tissues could determine which histological subtypes of their tumors could be developed and the patient prognosis as well.

**Figure 5.**
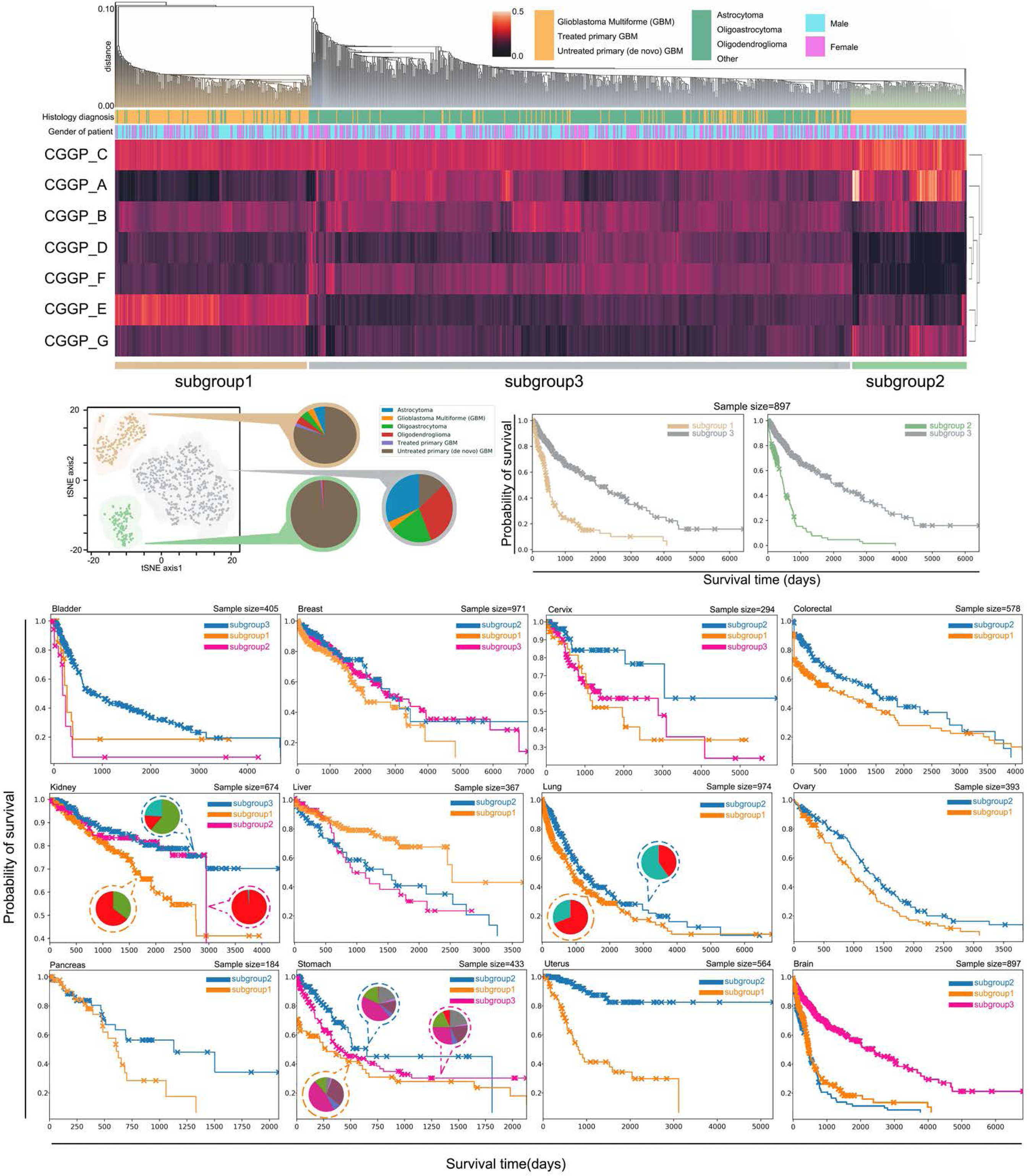
CGGP-defined subgroups were associated with cancer histological subtypes, oncogenic pathways, and clinical outcomes. (**A1**) Brain tumor patients were partitioned into subgroups with distinct clinical and biological tratis through unsupervised hierarchical clustering of the distribution profile of genomic patterns in patients’ germlines. (**A2**) Visualization of t-SNE illustrated the subgroups identified in A1. (**A3**) Significantly different survival outcomes of the subgroups (p<10^−15^). (**B**) Significant survival differences between germline subgroups in bladder, breast, brain, cervix, colorectal, kidney, liver, lung, ovary, pancreas, stomach and uterus cancers. Furthermore, germline subgroups were associated with tumor histological subtypes were shown in brain, kidney and lung cancers. Cancer types with insufficient sample size or histology diagnosis information were excluded from analyzes.

In 12 common cancer types (bladder, brain, breast, cervix, colorectal, kidney, lung, ovary, pancreas, stomach, and uterus cancers), CGGP-defined germline subgroups were significantly associated with prognosis (p=5.5×10^−3^, 2.2×10^−31^, 2.9×10^−3^, 3.0×10^−2^, 6.5×10^−19^, 5.6×10^−4^, 1.1×10^−2^, 1.2×10^−2^, 4.3×10^−2^, 8.9×10^−8^, 6.3×10^−18^, respectively; for cancer types with subgroups more than 3, reporting the most significant p-value) (Figure 5B). These results suggested that for these 12 common cancer types, clinical outcomes were partially determined by the CGGPs in germline genomes.

These results implied that CGGP profiles in germlines could also have influences on the oncogenic pathways in tumor tissues. In each of the above 12 cancer types, we selected the differentially expressed genes between subgroups and performed functional enrichment analysis to examine the affected oncogenic pathways (see Methods). The main differences between subgroup 1 and 3 of brain cancer patients were the epidermal growth factor (EGF) and insulin-like growth factor (IGF) pathways, whilst subgroup 2 and 3 differed in isomerase and mitochondria functions (Figure 6, Supplementary Materials Data D3). Of note, subgroup 1 and 2 were both mainly consisted of GBM patients but diverse in carrier status of germline patterns. Pathways related to immune response and responding to virus infection were differentially expressed in tumor tissues between the subgroups of at least 7 cancer types. Differential expression of the biological processes related to cell cycle and DNA damage/repair between the subgroups in 6 cancer types. TNF pathway, Wnt pathway, T cell signaling pathway, interferon signaling pathway and cell-cell adhesion pathway were found differentially regulated in the subgroups of breast, cervix, colon, kidney, liver, lung and uterus cancers. Further, functions related to mitochondria and epigenetic regulation were also observed to be prevalent between CGGP-defined subgroups (Figure 6).

**Figure 6.**
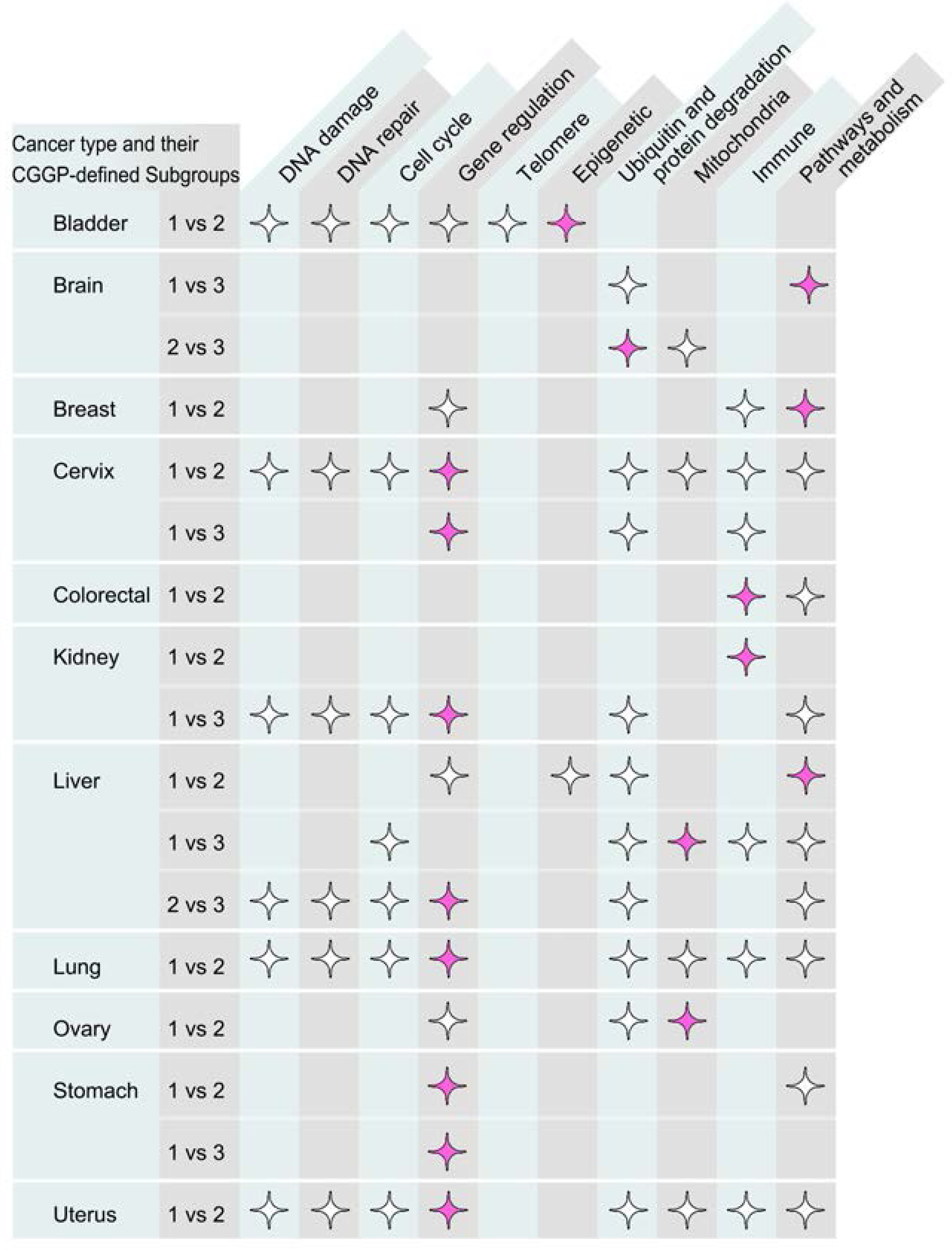
CGGP-defined subgroups showed significant differences in expression profile of corresponding tumor tissues. Stars marked that the functionality group had significant differential expression between the given patient subgroups. Colored stars stood for the most significant functionality groups for each subgroup pairs. For example, CGGP-defined subgroups of bladder cancer demonstrated differential gene expression profiles of functionalities including DNA damage & repair, cell cycle, gene regulation, telomere and epigenetic, with the last one being the most dictinct term.

Next, we examined whether normal tissues had different gene expression programs between the CGGP-defined subgroups. With a limited number of normal tissue expression profiles made available in TCGA, only cancer types with sufficient overall and subgroup-wise size were selected. Breast (n=76), colorectal (n=47), kidney (n=122) and lung (n=95) cancers were selected for this analysis. We identified differentially expressed genes in the normal tissues between the subgroups of breast, colorectal, kidney and lung cancers, respectively. Genes were functionally clustered and annotated using the same method described above. GO terms of biological processes, molecular functions and pathways showed significant differences between subgroups (Supplementary Material Data D4 and Table S3). For instance, differentially expressed genes between subgroup 1 and 3 of breast cancer were enriched in the following biological processes and functions: cell cycle (FDR-corrected p=2.0×10^−19^), mitosis (3.3×10^−18^), mitotic nuclear division (p=3.4×10^−14^), cell division (1.1×10^−13^), centromere (2.4×10^−9^) and others. The modulated genes between subgroup 2 and 3 of breast cancer have enriched functions and pathways which are related to ribosome and mRNA splicing differed (p<10^−10^). The modulated genes between the two subgroups of colorectal cancer, which also had survival differences, were enriched with transmembrane helix and region (p<10^−4^), which agreed with previous reports on relationship between transmembrane protein and colorectal cancer(Hashida et al., 2003). Interestingly, the above results implied that the impacts CGGPs imposed on patients’ tumor tissues were prominent but indirect; for example, in colorectal cancer, divergence in functions of transmembrane protein/region and signal peptide in normal tissue eventually exhibited differences in immune response functionalities in tumor tissue. Divergence of ATP-related metabolism between kidney cancer’s normal tissue subgroup1 and 2 transformed to differences in immune response functions in tumor as well. This was in accordance with the fact that carcinogenesis diseases are influenced by both genetics and environmental factors.

These results implied that contributions of CGGPs in patients’ germline genomes could play an important role in affecting on gene regulation programs in normal tissues, and then exert evo-constraints on oncogenic pathways in tumor progression and metastasis.

## Discussion

In this study, germline genomes paired with tumor genomes of 9,712 cancer patients and germline genomes of 16,670 non-cancer individuals were analyzed to reveal enriched sequential-context-dependent variant profiles which could be associated with cancer risk, tumorigenesis, and clinical outcomes. CGGPs provide an aspiring method for examining of the impact of germline variants on cancer development, latent molecular mechanisms and clinical outcomes. The interaction of genetic and environmental factors often leads to tumorigenesis. We showed that individuals with CGGP_E could be sensitive to tobacco smoke for developing 13 cancer types. Thus, for an individual who has a higher weight of CGGP_E in his/her germline, it could be possible to reduce the risk of cancer development through actively avoiding exposures to tobacco smoke. Surprisingly, although a higher contribution of the CGGP_E in germlines conferred a poor prognosis in various cancer types, such an association was not observed in lung cancer patients. One of the explanations could be that tobacco mutagens are directly exposing to lung so that tobacco mutagens could overpower the conditional genetic factors for lung cancer development and metastasis and thus, the clinical outcomes of lung cancer patients could be mainly driven by tobacco smoking. This hypothesis agrees with that fact that more than 85% of the lung cancer patients are smokers. On the other hand, other cancer types such as bladder cancer, the tissues are indirectly exposed to tobacco mutagens so that germline CGGP_E could play a more important role in tumorigenesis and outcomes.

Distinct combinations of CGGPs in germlines were associated with instinct cancer types, suggesting that germline genetic architecture with different fractions of the CGGPs has an impact on tumorigenesis in a tissue-specific manner, moreover, has evo-constraints on many aspects of the tumorigenesis and metastasis. For example, the germline subgroups defined by CGGPs were significantly associated with tumor histological subtypes, mechanisms of carcinogenesis and prognosis at least in 12 common cancer types, and the strengths of CGGPs in germline genomes are associated with key somatic cancer drivers in paired tumors. We also showed that CGGPs and their combinations in patients’ germlines have an evo-constraint on key somatic mutated genes and mutational processes (COSMIC signatures) in paired tumors. It seems that they are mainly associated with higher genome instability. In fact, germline genetic architecture could also impact on the gene regulatory programs in normal tissues. For example, we demonstrated that gene expression programs of the normal tissues between the CGGP-defined subgroups in at least 4 cancers are significantly modulated and enriched in cell cycle or other general biological processes which could contribute to cancer progression and metastasis. These results suggest that congenital germline variants encode some subtle causalities of tumorigenesis and metastasis. The quantitatively analyzing of germline variants provides novel perspectives of genomic features, which are associated with genetic diseases. We speculate that other genetic diseases could also be affected by their own specific germline genomic patterns. Furthermore, clustering analysis of germline genomes using the genomic patterns puts forward a novel notion of population scale genomic clustering, whereas previous efforts have been focusing on representing general geographic subgroups through gene-based evolutionary features(Rosenberg et al., 2002),(Tishkoff et al., 2009).

To quantify germline-defined intrinsic cancer risk, however, would require much larger cohorts that can represent the generic population, and more sophisticated evaluating approach for the weighing factors. In this study, we focused on the proof of concept, i.e. proving that the existence of germline genomic patterns, their impacts on somatic mutational events and clinical outcomes. Stricter criteria were chosen when we needed to translate consecutive weighing factors (in forms of float numbers) to binary CGGP assignments for these purposes. In theory, it is possible to assume a given individual to have a higher or lower risk of cancer by utilizing the CGGPs; but to accurately quantify intrinsic cancer risks, one needs more appropriate criteria. For example, in our stricter rules the boundary of confidence intervals was set to the point of 95% confidence, but it is widely acknowledged that the lifetime risk of cancer is around 30∼40% in the general population. This issue cannot be resolved simply by relaxing the boundary to 30% either, for we’ve showed that CGGPs are bettered considered in groups instead of independent individual patterns, and therefore, the intervals should be reconciled to this idea. Further investigations are needed to estimate intrinsic cancer risks in the future.

Finally, this study provides a conceptual advance in cancer genomics - the cancer-associated germline genomic pattern which is beyond the traditional cancer-risk genes. Genomic patterns could be a new type of genomic ‘dark matter’ encoded in germlines to influence cancer development and progression. They could also provide another dimension of genomic regulation for cancer development, progression and metastasis. Moreover, further investigation of germline genomic patterns could uncover genetic mechanisms of diseases that are beyond the gene-centered molecular mechanisms.

## Author contributions

E.W., Q.C. and X.F. designed the study. X.F. and X.X. performed data preparation, coding, mutational signature extraction and downstream analysis. D.L. contributed to examination of algorithms, distributed computing setup and feature selection methodologies.

## METHODS

Cancers from 22 primary sites were included in this study: adrenal gland, bladder, bone marrow, brain, breast, cervix, colorectal, eye, head and neck, kidney, liver, lung, lymph node, ovary, pancreas, pleura, prostate, skin, stomach, testis, thyroid, uterus. Note that although data derived from bone marrow, lymph node and thymus were collected and integrated into the mutational catalogs (see below), only solid tumor types were analyzed.

### Data and variant calling

To obtain germline variants of cancer patients, BAM files of the whole exome sequencing of buffy coats were collected from TCGA cohorts hosted at GDC Data Portal. The samples represent a non-redundant set of 9,712 individuals of 22 cancer types. Variant calling tool HaplotypeCaller from the GATK (version 3.8-0-ge9d806836; java -XX:ParallelGCThreads=4 - jar /thepath/GenomeAnalysisTK.jar -T HaplotypeCaller -nct 4 -R /thepath/GRCh38.d1.vd1.fa –I /thepath/bamfile.bam --genotyping_mode DISCOVERY -stand_call_conf 30 –o /thepath/bamfile.vcf) was used. We tested the HaplotypeCaller jointed calling mode and single sample calling mode by calling variants from Chromosome 1 of randomly selected 1,534 individuals from the 9,712 individuals, and found that the mutational catalogs derived from either jointed calling mode or the single sample calling mode highly resembled each other (Supplementary Materials Figure S4). Although samples representing blood cancers were included in the mutational catalog and the discovery of germline genomic patterns, corresponding individuals and cohort were not further analyzed due to the concern that tumor cells might contaminate the peripheral blood sample to the point undistinguishable for AF thresholding method. Considering the robustness of our methodology (Supplementary Materials), removing these samples in the initial stages would not significantly change the results reported in this study. To obtain the tumor somatic variants for the 9,712 patients, we retrieved their variants which were called by VarScan 2 and provided by TCGA (current release at GDC, v12.0; Supplementary Materials).

Control dataset, or the non-cancer dataset, consisted of non-cancer individuals collected from three cohorts: (a) Swedish Schizophrenia Population-Based Case-control Exome Sequencing (dbGaP ID: phs000473.v2.p2), representing 12,380 individuals of age 18-65. This cohort was also utilized by Genovese, Giulio, et al(Genovese et al., 2014) for investigating germline and hematopoiesis-derived somatic mutations. (b) Myocardial Infarction Genetics Exome Sequencing Consortium (dbGaP ID: phs001000.v1.p1) collected the whole-exome sequencing data of 2,322 non-cancer individuals. (c) Myocardial Infarction Genetics Exome Sequencing Consortium: Ottawa Heart Study (dbGaP ID: phs000806.v1.p1) contains whole-exome sequencing data of 1,968 non-cancer individuals. No individual was shared by the three cohorts.

### Criteria of determining germline variants

Only variants with read depth (DP) no less than 20 were retained for further inspection. Theoretically, germline homozygous and heterozygous variants would have VAF signals reside around 1.0 and 0.5, respectively, with subtle influences introduced by the systematic sequencing and variant calling bias. Among the variants derived from peripheral bloods of cancer patients, we observed a visible subset with VAF ranged around 0.2∼0.3, implying that putative somatic mutations generated from clonal hematopoiesis existed across the population (Supplementary Materials Figure S2). To avoid picking up somatic variants emerged from clonal hematopoiesis, which was a readily detectable process(Xie et al., 2014), (Jaiswal et al., 2014) in elderly people (>65 years old, 10%) and young population (1%)(Genovese et al., 2014), we developed a method which is similar to that described by Genovese et al.(Genovese et al., 2014) where through fitting a Gaussian mixture model (GMM) on the distribution of VAFs, we were able to determine the threshold for the germline variants. In TCGA cohort, GMM component revealed that 95% confidence interval of VAF for heterozygous variants was 0.422∼0.540. This interval was close to that reported previously (Genovese et al., 2014). Interval of non-cancer dataset was calculated independent from patient germline and yielded a similar scope (0.420 to 0.537). Criterion of VAF>0.9 was imposed for the homozygous variants, as we observed very few variants fell into the interval of 0.5∼1.0 and GMM model could not fit well around 0.9∼1.0.

### Discovering of genomic patterns using the non-negative matrix factorization method

We termed ‘trinucleotide profile’ of a variant as the trinucleotide form by the immediate 3’ sequential context of the variant (i.e., the previous nucleotide), the variant itself, and the immediate 5’ sequential context (i.e., the next nucleotide). Previous studies in tumor somatic-mutational signatures tended to focus on transcribed strand only because of strand bias. We, however, decided to include both transcribed strand and nontranscribed strand to achieve presumably better signal capturing. In this study, substitutions have 12 possibilities and each of the two context slots has 4 choices, which buildup 192 potential trinucleotide profiles. For each population (i.e., cancer germlines, cancer somatic mutations and non-cancer germlines), the corresponding mutational catalog was a 2D matrix of shape (192, number_of_samples), where each row was filled with non-negative integers that counted the number of presences of the given profile within a sample.

Non-negative factorization (NMF) factorizes a matrix V_i×j_ into two smaller matrices, W_i×k_ and H_k×j_. We modified the method developed previously(Alexandrov et al., 2013b) by introducing a hyperparameter that encouraged the exploration of distinct patterns through penalties addressed the unsparseness of pattern matrix. The rationale behind this was that germline genome, by nature, carried much more variants compared to somatic mutations in tumor genome, and the variants were less informative than the *de novo* mutations in tumors (i.e., somatic mutations in tumors are more closely associated with tumorigenesis, whereas germline variants are by defination inherited from parents, and do not directly associated with specific biological process or disease in most cases, thus ‘noisier’). The number of germline genomic patterns was determined using the silhouette method, which was previously implemented for extracting tumor mutational signatures(Nik-Zainal et al., 2012). The silhouette score (i.e. the stability of the solved genomic pattern) would gradually decayed as more germline genomic patterns being extracted (k=2 to k=7), and then drop drastically from k=7 to k=8, k=8 to k=9 and so on, where the score decreased from more than 0.5 immediately to 0.2 or less. Also, in k>7 setups, the extra patterns would closely resemble previously known patterns (i.e. the ones found by k=7 setup; cosine similarity<10^−5^). The k=7 was the last point before overfitting. We tested the robustness of this NMF approach in several ways, where we showed that the approach was able to tolerate random noise. For example, randomly adding or removing of up to 30% noise to the mutational catalog allowed the reproducing of original germline genomic patterns (see Supplementary Materials for details). Germline genomic patterns, once made available, could be assigned ubiquitously to given germline genomes through converging NMF model without updating the pattern matrix (W_ixk_). In other words, new individuals could be evaluated by the germline genomic patterns easily.

### Clustering and visualization of the CGGP-defined patient subgroups

Hierarchical clustering was conducted based on the Euclidean distance and the Ward’s method. Similar results were obtained with or without PCA prior to clustering. The tSNE(van der Maaten and Hinton, 2008) is a dimension reduction and visualization algorithm that has been applied to various fields including GWAS (MejŁa-Roa, 2015) and protein similarity comparisons (Ray, 2016). This algorithm provides meaningful visualization of latent clusters inside high dimension data, which could serve as an aid to view hierarchical clustering results more intuitively.

### Examining of CGGPs’ impact on COSMIC somatic signatures and important somatic mutations in ontogenetic genes

We assigned COSMIC somatic signatures using the method and criteria reported in their publication{Alexandrov:2013eb}(Supplementary Materials). For each CGGP, patients of each cancer type were then sorted in descending order of weighing factors, and individuals in the top/bottom quantiles were selected to form to subgroups. Fisher’s test was used to examine whether higher weight of the CGGP in germline would imply significantly more possession of a COSMIC signature.

For a given gene, patients of each cancer type were partitioned based on their somatic mutation status of the gene, which were either determined by TCGA’s clinical records or variant calling results. Weighing factors of CGGPs were compared between the two subgroups through t-test to examine whether enrichment of any of the germline genomic patterns were associated with enrichment of important somatic mutations in tumor tissue. Using binary CGGP assignment (see Supplementary material) and Fisher’s test would give similar results.

### Functional enrichment analysis of differentially expressed genes between the CGGP- defined caner subgroups

Same method was used for germline data and somatic data. For individuals in a given tumor type and sample type (i.e. somatic or germline), we obtained their gene expression profiles (raw counts) from GDC repository. Multiple probe reads for a given gene were averaged. We ranked all probed genes based on their level of differential expression, measured by t-test, between the two germline-defined subgroups, and selected either the most significant 3000 genes or all genes that had p-values less than 1.0×10^−3^. Raw counts and unadjusted p-values were used because our purpose here was to rank genes, instead of a formal differential expression analysis. We fed the picked genes into Functional Annotation of DAVID 6.8 for functional annotation and clustering.

### Data and Code Availability

All data generated or analysed during this study are included in this article and supplementary information files.

